# Model mimicry limits conclusions about neural tuning and can mistakenly imply unlikely priors

**DOI:** 10.1101/2024.01.31.578040

**Authors:** Michael J. Wolff, Rosanne L. Rademaker

## Abstract

In a recent issue of Nature Communications, Harrison, Bays, and Rideaux^1^ use electroencephalography (EEG) to infer population tuning properties from human visual cortex, and deliver a major update to existing knowledge about the most elemental building block of visual perception – orientation tuning. Using EEG together with simulations in an approach they refer to as “generative forward modeling”, the authors adjudicate between two competing population tuning schemes for orientation tuning in visual cortex. They claim that a redistribution of orientation tuning curves can explain their observed pattern of EEG results, and that this tuning scheme embeds a prior of natural image statistics that exhibits a previously undiscovered anisotropy between vertical and horizontal orientations. If correct, this approach could become widely used to find unique neural coding solutions to population response data (e.g., from EEG) and to yield a “true” population tuning scheme deemed generalizable to other instances. However, here we identify major flaws that invalidate the promise of this approach, which we argue should not be used at all. First, we will examine the premise of Harrison and colleagues^1^, to subsequently explain why “generative forward modeling” cannot circumvent model mimicry pitfalls and can deliver many possible solutions of unknowable correctness. Finally, we show a tentative alternative explanation for the data.

**Conflict of interest:** The authors declare no conflict of interest

## Main Text

Invasive neural recording techniques are the gold standard and only direct measurement tool for quantifying neural orientation tuning properties in visual cortex. Harrison and colleagues^1^ point to previous research as precedence for the overrepresentation of horizontal compared to vertical selective neurons in visual areas of mouse^2^, cat^3^, and macaque^4^, and summarize these three studies in Figure 1b of their paper^a^. However, Harrison and colleagues seem to misinterpret data from these studies that do not, in fact, set out to test for or convincingly show anisotropies between vertical and horizontal orientations. Roth and collegues^2^ show a weak trend of more horizontal versus vertical selectivity in mouse V1, but an *opposite* trend in a later visual area (Posteromedial area). Wang and collegues^3^ show a similar weak trend favoring horizontal selectivity in cat visual cortex, but other cat studies that show no or *opposite* trends can be easily found (e.g., ^5,6^). Fang and colleageus^4^ analyzed data from a total of 48 V1 hemispheres and 38 V4 hemispheres (in over 34 macaques), showing no trends favoring horizontal orientations in V4^b^, and an *opposite* trend in V1^c^. Thus, even this small selection of research from a field that spans decades (starting with Hubel & Wiesel in 1962^7^) shows no consistent differences between vertical and horizontal neural selectivity, and it is unclear to us how Harrison and colleagues can argue otherwise. If systematic anisotropies between vertical and horizontal selectivity exist, we are not aware of research that has systematically evaluated the existing literature or actually applied quantitative tests. Furthermore, the implied evolutionary justification for an anisotropy favoring horizontal over vertical orientations, seemingly mirrored in the statistics of natural but not man-made scenes^d^, is difficult to reconcile with decades of physiological research emphasizing developmental influences shaping orientation tuning^8–10^.

**Figure 1.**
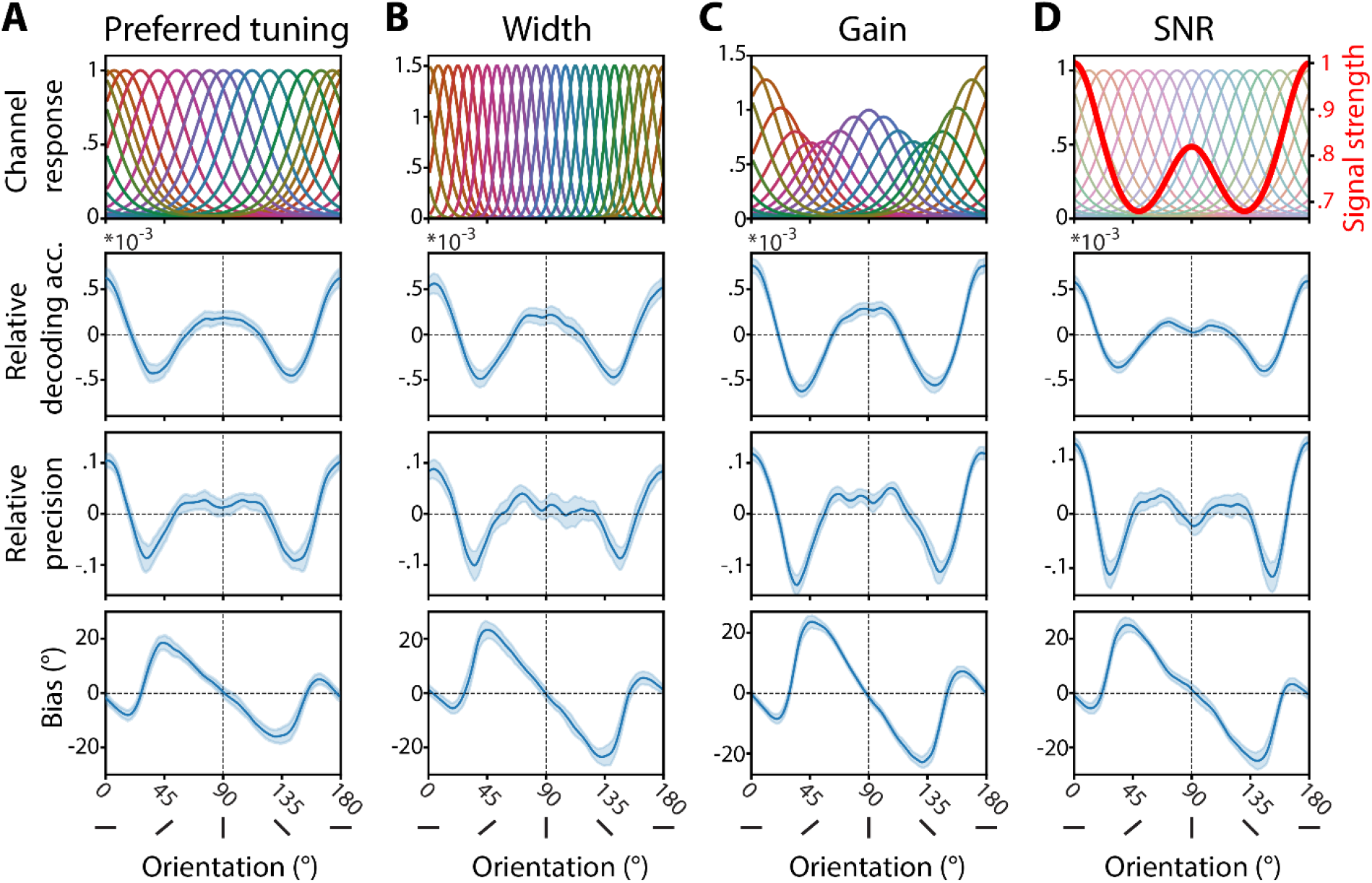
Model mimicry: many models produce the same pattern of results. In “generative forward modeling”, EEG data are simulated from models that use different sets of orientation tuning functions (top row). Decoding results (relative decoding accuracy, relative precision, and bias) as a function of orientation are shown (3 bottom rows) for simulations using different underlying example models. **A**. Preferred tuning model: Tuning functions are unevenly spaced along the orientation space, with more clustering at vertical, and even more at horizontal orientations. This is the “best fitting” model from Harrison and colleagues^1^. **B**. Width model: Tuning curve widths are uneven, with narrowest tuning for obliques, wider tuning for vertical and widest tuning for horizontal. **C**. Gain model: Uneven tuning curve gain across orientations space, with more gain at cardinals that is highest for horizontal orientations. **D**. SNR model: Tuning curves are uniform, but signal strength is orientation specific.

Aside from its shaky premise, the central flaw in Harrison and collegues^1^ lies with the fact that EEG decoding results cannot inform about the underlying neural or population tuning, due to an inherent inverse problem and model mimicry. The inverse problem is where an underlying cause cannot be inferred from a (measurable) effect, such as the inability to estimate neural causes from non-invasive imaging results^11,12^. Relatedly, model mimicry refers to cases where many possible models can generate the same, or very similar, outcomes and model fits. To test what population tuning properties explain their pattern of EEG results, Harrison and colleagues^1^ claim they can use “generative forward modeling” (see also^13^) to differentiate between two possible population tuning schemes: differences in tuning widths and differences in tuning preferences. This claim is false. Using the same simulation approach as Harrison and colleagues (but a slightly different decoder^e^), we show some examples of population tuning schemes that all yield the same pattern of results at the macro-level (Figure 1, Supplemental Methods). Importantly, this includes the population tuning scheme with different tuning widths that Harrison and colleagues argued could not fit their data. This is because they did not consider a wider parameter space for this model. Even when adjudicating between just two models, there are many possibly “sensible”, but fairly arbitrary parameters to pick from (e.g., the number of tuning functions, the range of tuning function widths, etc.) that can all generate outcomes that mimic each other. Excluding parts of this parameter space (such as models with wider tuning at cardinals compared to obliques) on the basis of previous physiology findings is not an option, as any subsequent claims about orientation tuning would amount to reverse inference. In addition to models with differences in tuning width or preference, a model with differences in gain modulation^14^ could also explain the data (Figure 1c), but was not considered by Harrison and colleagues^1^. In fact, even a uniform set of tuning curves can approximate the data, as long as the signal-to-noise ratio (SNR) is modulated across orientation space (Figure 1d). To make matters even more complex, model specification is not limited to “sensible” choices only – the “tuning functions” used by Harrison and colleques^1^ are well-motivated models^15^, but a set of arbitrarily shaped functions could also be used to simulate data and/or recover decoding metrics^16^.

Across two EEG data sets, Harrison and colleagues^1^ show that horizontal orientations result in notably better and more consistent decoding than vertical orientations. Given that “generative forward modeling” cannot be used to infer orientation tuning anisotropies in human visual cortex as a plausible explanation, what other factors might be driving these EEG results? Using the same decoding method as for the simulations (Figure 1), we replicate the pattern of results both for the openly available data^13^ used by Harrison and colleagues^1^, as well as for another openly available EEG data-set where orientated grating stimuli were presented centrally^17^ (Figure 2A). However, this effect is not replicated for EEG data sets^18,19^ where orientation gratings were presented laterally (Figure 2B), where no differences between vertical and horizontal orientations are evident.

**Figure 2.**
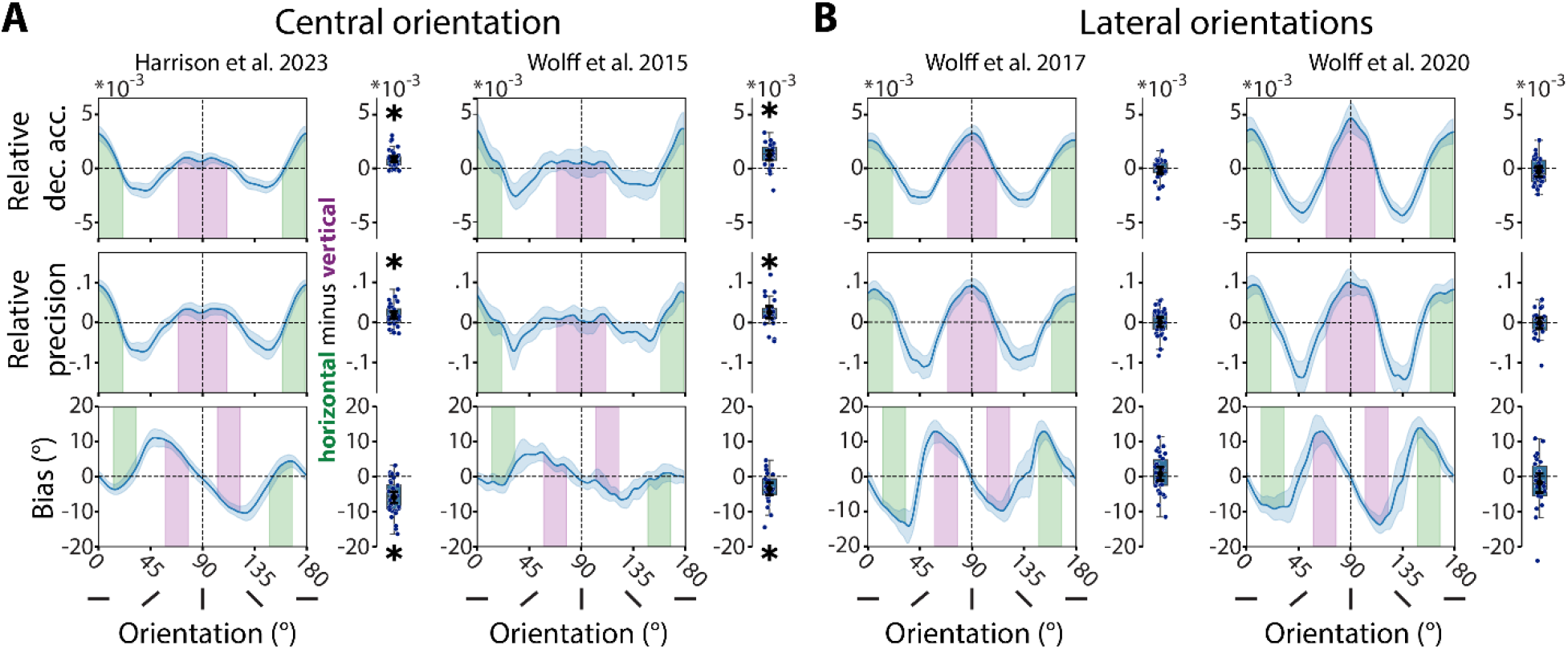
Cardinal anisotropies for orientation decoding are specific to central stimuli in EEG. **A**. Re-analyses of experiments reported by Harrison and colleagues^1^ (*left*) and Wolff and colleagues^17^ (*right*) where central orientations were shown to participants. Line plots show relative Mahalanobis distance-based decoding metrics as a function of orientation, with shaded areas indicating the cardinal orientation bins used to compute differences between horizontal (green) and vertical (purple) orientations. Box plots show decoding metric differences for horizontal minus vertical orientations. *Top*: Relative accuracy (mean-centered cosine vector mean of pattern similarity curve), *Middle*: Relative precision (1 minus the circular standard deviation of decoded orientation across trials), *Bottom*: Bias of pattern similarity curves, in degrees. Error bars are 95% C.I. Both data-sets show consistent differences between horizontal and vertical orientations (^*^ *p* < 0.05), with better decoding for horizontal orientations, and a stronger attraction bias toward vertical orientations. **B**. Re-analyses of experiments with laterally presented orientations^18,19^. Same conventions as in A. No consistent differences between horizontal and vertical orientations.

Why do we see this difference between centrally and laterally presented stimuli? We hypothesize that the cardinal anisotropy seen only for centrally presented gratings is driven by visual field anisotropies – i.e., anisotropies of location instead of orientation. Human visual performance is higher for stimuli presented along the horizontal compared to the vertical meridian, especially peripherally, and human V1 has about double the cortical surface area dedicated to the horizontal compared to the vertical meridian^20,21^. In EEG, further anisotropies may arise due to the organization of the visual field map in cortex, which determines how well activity from different portions of cortex are captured by EEG scalp electrodes. For example, locations along the vertical meridian are processed closer to, and inside of, the longitudinal fissure^22^, which is more difficult to measure with scalp electrodes.

That differences between the vertical and horizontal meridians of the visual field play a role in EEG measurements, becomes evident when looking at location decoding from these signals. We re-analyzed multiple openly available EEG datasets where participants were presented with a single dot at one of many possible locations around fixation^23–25^. We see clear and systematic differences in location decoding accuracies, with highest relative decoding for locations close to the horizontal meridian, and lowest for locations close to the vertical meridian (Figure 3a). The eccentricity at which dot stimuli were presented in four of these datasets (outer ring in Figure 3a, *bottom*) overlaps with the edge of the grating stimulus used by Harrison and colleagues (overlaid grey dotted line in Figure 3a, *bottom*)^f^. Concretely, for centrally presented stimuli, this difference in sensitivity across the visual field means lower SNR at the upper and lower stimulus edges than at the left and right stimulus edges (Figure 3b, *left*).

**Figure 3.**
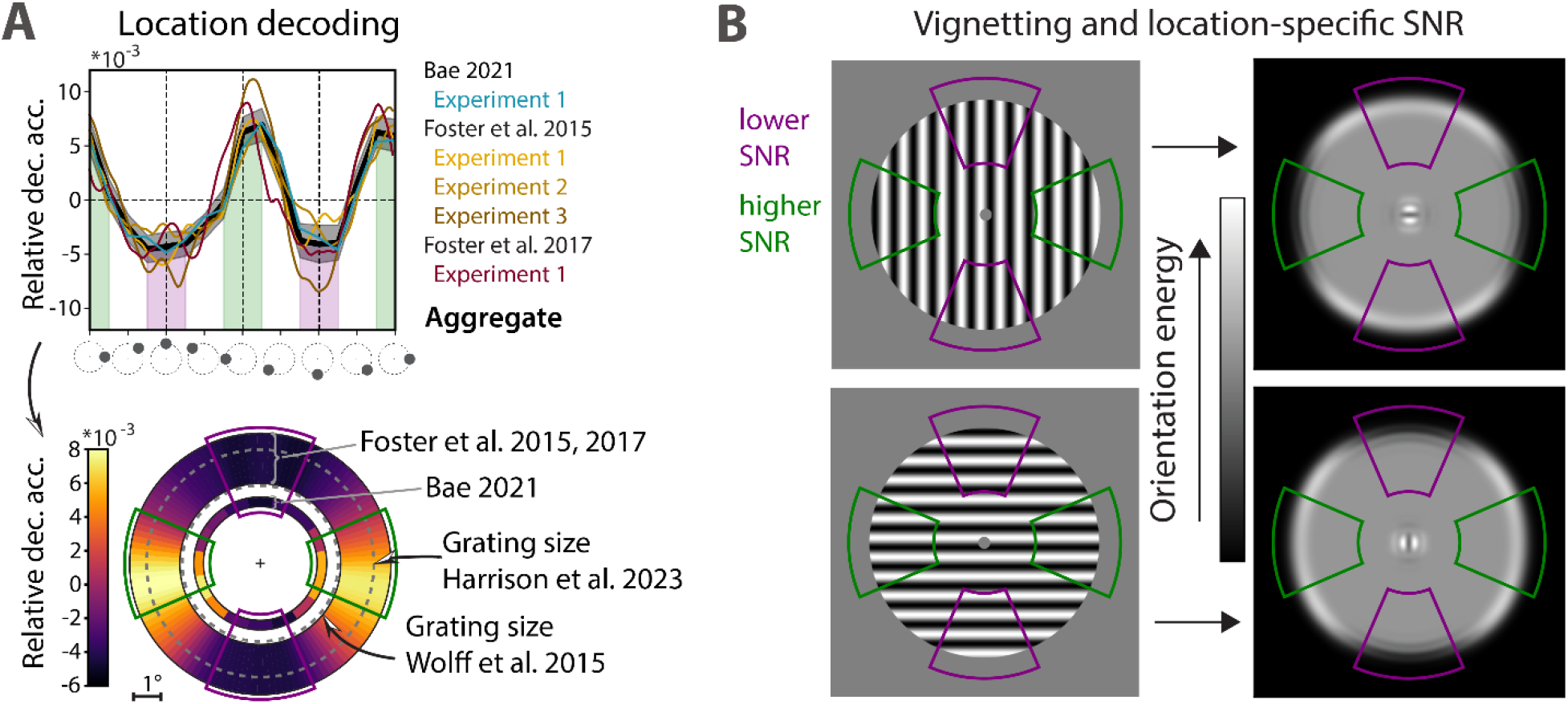
Differences in measurement sensitivity across the visual field that interact with stimulus vignetting can explain decoding differences between vertical and horizontal orientations that are presented centrally. **A**. *Top*: Relative location decoding accuracy as a function of presented stimulus location (re-analyses of ^23–25^). Green and purple shadings highlight stimuli presented on the horizontal and vertical meridians, respectively. *Bottom*: Location decoding across the visual field, plotted to scale for the various experiments: Outer, thicker ring represents possible locations of dot stimuli used in Foster and colleagues^24,25^, presented at 3.8°–4° eccentricity and 1.6° in diameter. Inner ring represents possible locations of the dot stimuli used in Bae^23^, presented at 2.3° eccentricity with 0.35° diameter. Dashed grey circles represent stimulus sizes of the central orientations used in Wolff and colleagues^17^ and Harrison colleagues^1^ (radii of 2.88° and 4.2°, respectively). **B**. Vignetting^26^ for gratings presented at the center of the screen: “Orientation energy” is highest on the stimulus edges aligned with the orientation. This means relatively higher orientation energy along the vertical meridian for vertical orientations (*top)* where SNR is low, and along the horizontal meridian for horizontal orientations *(bottom)* where SNR high.

Importantly, these SNR differences can interact with second order stimulus properties (stimulus edge effects or “vignetting”) that have been argued to at least in part be related to the decoded signal obtained from non-invasive neuroimaging^26,27^. Vignetting refers to the interaction between stimulus orientation and stimulus aperture, such that for circular gratings the “orientation energy” is strongest on the edges of the grating aligned with the orientation (Figure 3b, *right*). A vertically orientated grating presented centrally will therefore evoke more activity in the periphery of the vertical meridian, a visual field location where sensitivity is lower. A centrally presented horizontal grating will evoke more activity in the periphery of the horizontal meridian, where sensitivity is higher. This may not be true for laterally presented stimuli, where the decoded orientation energy falls into a part of the visual field where measurement sensitivity is more evenly distributed.

We do not claim that this explanation is definitive or exhaustive. For example, spatial attention to the endpoints of orientated gratings^28^ could interact with visual field anisotropies in a manner similar to vignetting effects. Other factors may also interact with measurement of orientation selectivity, such as stimulus contrast^29^ or radial bias^4,30^. We want to highlight the importance of considering stimulus and measurement biases that can interact with orientation decoding. Finally, the well-known “oblique effect” which describes better perceptual performance for cardinal over oblique orientations^31^ and is mirrored in the overrepresentation of orientation-tuned neurons that prefer cardinal over obliques^2–4^, aligns with the EEG results for both centrally and laterally presented gratings (see Figure 2). This implies that at least some form of orientation anisotropy may be genuinely measurable with EEG. That said, we argue that the observed attenuation of decoding metrics for vertical compared to horizontal orientations, specific for centrally presented gratings, is likely driven by second-order stimulus properties that interact with location-specific measurement differences.

In conclusion: Despite decades of research, invasive neural recordings in animals have not found the anisotropies between vertical and horizontal orientations seen in the EEG data reported by Harrison and colleagues^1^. This pattern of results cannot be explained on the basis of the underlying orientation tuning, because “generative forward modeling”^1,13^ suffers from an inherent inverse problem, where many possible population tunings can approximate the patterns of reported EEG data equally well. Given that the pattern of EEG results does not replicate for laterally presented stimuli, cardinal anisotropies are likely driven by other factors, such as differences in visual field sensitivity between the vertical and horizontal meridian and their interaction with second-order stimulus effects.

## Acknowledgements

We want to thank the multiple groups of scientists whose work we re-analyzed here. We were able to test our alternative hypotheses thanks to their efforts of ensuring readily available and well-documented data. We’d also like to thank Tommy Sprague and John Serences for reading an early version of this manuscript, and for their valuable feedback.

## Supplemental Methods Simulations (generative forward modelling)

We simulated different population tuning models using largely the same approach as in Harrison and colleagues^1^ by adapting their published Matlab scripts. Briefly, data for 36 “ subjects”, from 32 “ EEG channels” and 6480 trials per “subject”, was simulated for each model (see below). Each model had a given number of tuning functions, or “model channels”. The modelled channel responses to each orientation (1° to 180°, in steps of 1°) “shown” to a given model were transformed to EEG sensor space via matrix multiplication between the orientation-specific model response of each trial, and a random weights matrix (number of channel functions by number of “ EEG channels”, sampled from a uniform distribution over 0 to 1). Trial-specific noise was added to each simulated “ EEG channel” sampled from a normal distribution (s.d. = 6), which also ensures differences in simulated responses to trials on which identical orientations were “shown”.

For all models we will only mention any deviations from Harrison and colleagues^1^ “Preferred tuning model”. The purpose of our simulations was to demonstrate that there is no unique model that best describes the data, even when only considering models that could be argued to be plausible. Our models (Figure 1) are by no means the “best fitting” models, as searching for best fitting solutions in this very large parameter space would be computationally intractable. Indeed, we derived at our models through mere trial and error and stopped once we obtained decoding results that resembled Harrison and colleagues^1^ “Preferred tuning model”. Thus, the models and their parameters described below should not be considered “definitive”; they are snapshots out of many more possibilities.

Preferred tuning model: For this we used the exact script published by Harrison and colleagues^1^, which generates the preferred tuning model. This model consisted of 16 tuning functions with constant widths (κ = 2). Preference was modulated by shifting the tuning functions based on the sum of two von Mises derivative functions (κ = 0.5) centered on 0° (amplitude = 14) and on 90° (amplitude = 8), which has the effect that there are relatively more tuning functions around horizontal (0°) compared to vertical (90°), and fewest tuning functions around obliques (45° and 135°). Note that these values in the scripts uploaded by the original authors at the time of writing, differ slightly from the values described in the manuscript (which states the amplitudes were 15 and 10). The resulting difference between these two parameter settings is marginal however, and we decided to stick to those parameters in the script as uploaded by the authors, without changing anything.

Width model: Instead of changing the tuning preferences across the orientation space, tuning functions were evenly spaced, but their widths were modulated. This modulation was derived from the *inverse* of the sum of two von Mises functions (κ = 0.5), one centered on 0° (amplitude = 15) and one centered on 90° (amplitude = 4). Given the inversion, tuning widths were wider for cardinals than for obliques, with horizontal widths being wider than vertical widths. The possible tuning widths were rescaled such that they ranged from κ = 7 (the widest) to κ = 19 (the narrowest). The number of tuning functions were increased to 24 (from the original 16) and every tuning function were scaled to range from 0 to 1.5.

Gain model: The gain model comprised 16 evenly spaced tuning functions with constant widths (κ = 2), but differences in scaled amplitude (“ gain”). Gains were modulated from the sum of two von Mises functions (κ = 0.5), one centered on 0° (amplitude = 15) and the other on 90° (amplitude = 8). The range of gains were scaled from 0.7 (at the obliques) to 1.4 (at 0 degrees, which is horizontal). SNR model: Here we used a uniform distribution of 16 identical tuning functions, all with the same width (κ = 2) and all with the same gain (amplitude of 1). Unlike the models above, here we do not manipulate the underlying tuning response functions, but instead modify the signal strengths of the simulated activity patterns across the “ EEG channels” . Specifically, the signal strength was modulated for simulated response patterns generated from each of the 180 orientations using the sum of two von Mises functions (κ = 0.5), centered on 0 degrees (amplitude = 15) and on 90 (amplitude = 8). Signal strength modulation ranged from 0.68 (68% signal strength) to 1 (100% of signal strength). The signal strengths of the orientation-specific patterns were modulated *after* transforming activations from the tuning response functions to every possible orientation (1°–180°) into sensor space (as described in Harrison and collegues^1^ and above), meaning that the orientation-specific patterns of the simulated EEG sensors were multiplied by the corresponding signal strengths (0.68 to 1), before adding the same amount of Gaussian noise to each (s.d. = 6).

## EEG Data

We reanalyzed openly available EEG datasets of 9 experiments across 7 publications ^1,17–19,23–25^, where human participants viewed either circular orientation gratings, or locations. For the present manuscript, the stimulus sizes and locations that participants viewed while EEG was recorded are of particular interest, and are described in more detail below. Other details are available in the methods sections of the original publications.

<u>Harrison et al. (2023)</u>^1^: Participants (*N* = 36) viewed serially presented, randomly orientated circular gratings (4.2° radius) centered around fixation. Each grating was presented for 50ms, with an ISI of 150ms between consecutive gratings. The task was to detect gratings with a lower spatial frequency.

<u>Wolff et al. (2015)</u>^17^: Participants (*N* = 24) performed a visual working memory task, where the orientation of a grating had to be memorized for up to 2.6 seconds. Each circular grating was centrally presented (2.88° radius) for 200ms, followed by a blank delay of at least 1.17 seconds.

<u>Wolff et al. (2017)</u>^18^: Only experiment 1 was reanalyzed. Here, participants (*N* = 30) performed a retro-cue visual working memory task. Two randomly orientated circular gratings (radius of 3.345° each) were simultaneously presented on the horizontal meridian at 6.69° eccentricity. The presentation time was 250 ms, followed by a blank delay of 800ms. The orientations of both gratings were behaviorally relevant during encoding.

<u>Wolff et al. (2020)</u>^19^: Participants (*N* = 26) also performed a retro-cue visual working memory task with laterally presented, randomly orientated circular gratings. The gratings (radius of 4.255° each) were presented at 6.08° eccentricity for 200 ms followed by a blank delay of 400 ms. The orientations of both gratings were behaviorally relevant during encoding.

<u>Foster et al. (2015)</u>^24^: Participants performed a spatial working memory task in all three experiments. The visual stimulus on each trial in all three experiments was a dark gray circle (0.8° radius) presented on a random location of an invisible circle at 4° eccentricity. The participants’ task was to memorize the location for a delay of at least 1s (variable across experiments). In experiment 1 and 3, the circle was presented for 250 ms and in experiment 2 for 1s. Sample size was N *=* 15 in all experiments.

<u>Foster et al. (2017)</u>^25^: We reanalyzed experiment 1 (*N* = 10). Here, in each trial a single a randomly colored circle (0.8° radius) was presented on a random location of an invisible circle at 3.8° eccentricity. Stimulus duration was 100ms, followed by a 1.2s blank delay. The participants’ task was to memorize and report the color of the colored circle in each trial.

<u>Bae (2021)</u>^23^: We reanalyzed experiment 1, where participants (*N* = 22), performed a spatial working memory task. The visual stimulus was a small circle (0.175° radius) presented for 200ms on one of 16 discrete locations on an invisible circle at 2.3° eccentricity. A blank delay (1.3s) followed after the offset of the circle. The task was memorize and report the location of the circle on each trial.

## Preprocessing

For all experiments, we used the voltage data the way it was published and preprocessed by the original authors.

For the subsequent decoding analyses, we used the voltage traces from 50 to 450 ms relative to stimulus onsets from the posterior electrodes, in line with Harrison and colleagues^1^. Data from ref^17–19^ and the reanalyzed experiment 1 from ref^23^ all used the same electrode coverage, and the same 17 posterior channels were included in the corresponding analyses (P7, P5, P3, P1, Pz, P4, P6, P8, PO7, PO3, POz, PO4, PO8, O1, Oz and O2). The same electrodes were included for the data of ref^1^ in addition to the electrodes Iz, P9, and P10. The electrode coverage was lower for the reanalyzed experiments in ref^24,25^ and the included posterior electrodes for these data-sets were PO3, PO4, P3, P4, O1, O2, POz, and Pz.

Instead of decoding at each time-point separately within the time-window of interest and then averaging (as in Harrison and collegues^1^), we first reformatted the data in a manner similar to previous work^19^ before feeding it to the decoder: To take advantage of the fact that stimulus-specific information is not only present in the activity patterns across electrodes, but also in the *temporal* pattern of the evoked voltage changes, we combined the channel and temporal dimensions to improve the sensitivity of the decoder. To do so, we first down-sampled the signal from the time-window of interest (50 ms to 450 ms, relative to stimulus onset) to 50hz (51.2 Hz for Harrison and collegues^1^, due to the original sampling rate of 1024Hz), and removed the mean activity level within each trial and electrode. The resulting, mean-centered 20 voltage values of each channel in each trial were then combined with the channel dimension. The number of dimensions for the decoder increased therefore 20-fold (number of down-sampled time-points by number of channels).

## Stimulus decoding

We used a Mahalanobis distance based decoder^19^ to decode orientations from the simulated data and orientations/locations from the spatiotemporal signals from the EEG data-sets. The approach was identical for both orientation and location decoding apart from taking into account that orientations are in 180° space, while locations are in 360° space. We used an 8-fold cross-validation approach. First, trials were assigned to the closest of 16 evenly spaced orientations/locations (variable, see below). The trials were then randomly split into 8 folds using stratified sampling. The trials of 1 fold were held out for “testing” and the trials of the remaining 7 folds were part of the “training data” . The covariance of the train trials was estimated using a shrinkage estimator^32^, before the number of trials in each orientation/location bin of the train data was equalized through random subsampling. The subsampled trials within each bin of the training set were then averaged. And the averaged bins were then convolved with a half cosine basis set raised to the 15^th^ power^33^ to pool information across similar orientations/locations. The Mahalanobis distances between the left-out test trials and the averaged train bins were then computed. This procedure was repeated for all train/test fold combinations. The experiment of one dataset^23^ used exactly 16 evenly spaced locations. Here the original location labels were used, rendering the aforementioned binning unnecessary. All remaining datasets used random orientations/locations, for which the above procedure was run separately for 8 possible ways of binning the orientations (with bins centered at 0° to 168.75°, at 1.40625° to 170.1563°, at 2.8125° to 171.5625°, at 4.2188° to 172.9688°, at 5.625° to 174.375°, at 7.0313° to 175.7813°, at 8.4375° to 177.1875°, or at 9.8438° to 178.5938°, each in 16 steps of 11.25) or location spaces (same as for orientation, but converted to 360° space by multiplying all values by two). This means that for each trial, we obtained 16 times 8 = 128 Mahalanobis distance bins (with the exception of the data-set with only 16 discrete orientation, which resulted in exactly 16 distances per trial). Given the randomness of the initial folds and the subsampling within folds, the above procedure was repeated 20 times to obtain more robust results. Once all distances were obtained and averaged over repetitions, distances for each trial were mean centered by subtracting the average distance across all Mahalanobis bins from each. The distances were then ordered as a function of angular difference between test and train bin, obtaining “pattern similarity curves” for each trial. For experiments with two simultaneously presented orientation gratings, one on each side^18,19^, each orientation was decoded separately.

### Relative decoding accuracy, relative precision, and bias

“Decoding accuracy” was obtained for each trial by computing the cosine vector mean of the “pattern similarity curve”^18^. “Decoding accuracy” was then averaged as a function of orientation/location using a sliding window (width = 11.25° for orientations, width = 25° for locations) that moved over angular space in steps of 1.40625°/2.8125° for orientations/locations. “Relative decoding accuracy” was obtained by mean-centering the resulting orientation/location –specific decoding accuracy curve. “ Precision” was obtained by taking the circular means of the trial-wise “ pattern similarity curves”, and calculating the inverse circular standard deviation over these means. “Relative precision” was obtained by mean-centering the precision curve (same as for “ relative decoding accuracy). “Bias” was obtained by computing the circular mean of the averaged “pattern similarity curves” of all trials within each angular window (same as above).

For visualization, the angular relative decoding, relative precision and bias curves were smoothed across orientations/locations with a Gaussian smoothing kernel (s.d. = 2°/4° for orientations/locations).

To explicitly test differences in decoding accuracy and precision between vertical and horizontal orientations (as shown in Figure 2), the respective decoding metrics were averaged from -22.5° to +22.5° relative to 0° and 90° degrees. For the bias we assumed that, given equal attraction towards each cardinal, the effect should be maximal for orientations 22.5° away from the cardinals, i.e., halfway the distance to the obliques, where the influence of each cardinal should be cancelled out. We thus averaged the bias values from -12.5° to 12.5° relative to 22.5° and 157.5° for horizontal orientations, and relative to 67.5° and 112.5° for vertical orientations, *after* sign reversing bias values such that positive values always correspond to attraction to the nearest cardinal.

## Edge effects (“vignetting”)

We used the “ perfect cube model“ ^26^ to illustrate a possible relationship between location-specific SNR differences, and “orientation energy”, strongest at the edges for circular gratings. We used the exact stimulus and model parameters as described in ref^26^. Briefly, two sine-wave gratings (one vertical, the other horizontal) were convolved with eight distinctly oriented 2D Gabor “filters” (0° to 157.5°, in steps of 22.5°), which all had the same spatial frequency as the sine-wave gratings. The output of each filter was normalized before taking the sum over all eight, resulting in the 2D “orientation energy” plot in Figure 3b.

## Statistical significance testing

The reported differences between horizontal and vertical orientations (Figure 2) were tested for significance using a permutation t-test with 10,000 permutation as implemented by the python toolbox MNE. All tests were two-sided and the statistical significance threshold was *p* < 0.05.

## Code availability

The code used to generate the figures and results reported in this manuscript are available at https://github.com/mijowolff/model-mimicry-and-unlikely-priors.

## Data availability

All data used in this study is openly available^17–19,23–25^. For convenience, data from^17–19^ was reduced in size by only including electrodes and time-points of interest, and is available at https://osf.io/bdf74/

---

**Supplemental figure 1.**
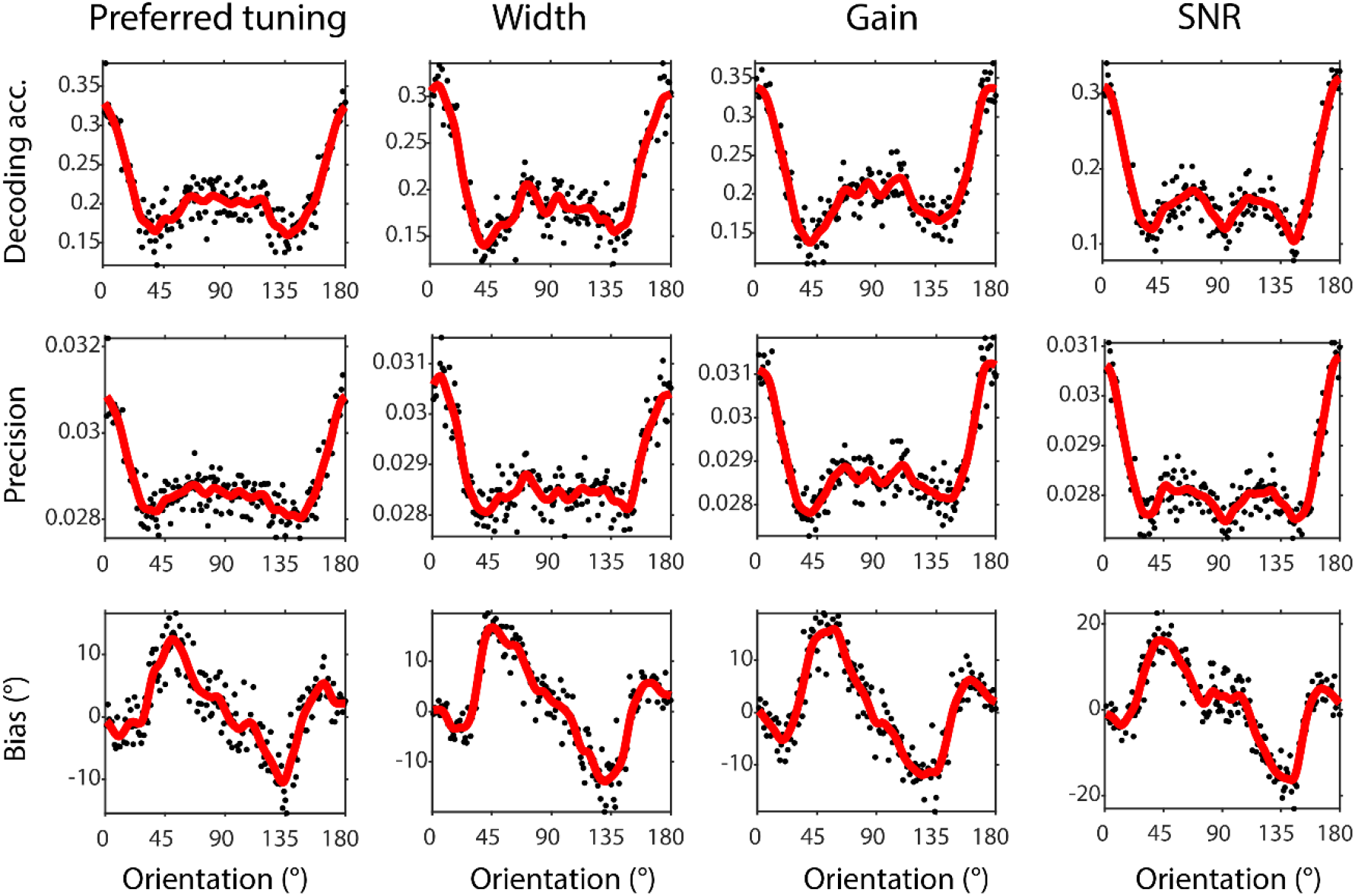
Same underlying models as in Figure 1, but the results and figures are obtained from Harrison and colleagues^1^ implementation of an IEM-based decoder and visualization approach by using their published Matlab code, but folded 10 times as described in their Methods.

## Footnotes

a How these data were derived and plotted is not described, and statistical tests of vertical-horizontal anisotropies are not reported.

b Incidentally, the first data figure in Fang and collegues^4^ shows a single example V4 hemisphere, and here a trend for more horizontal-preferring neurons can be observed. This trend is absent in the full V4 data with 38 hemispheres.

c This higher selectivity for vertical is statistically significant, but likely due to concurrent radial biases.

d The justification for an embedded prior based on natural scene statistics (i.e., the green line in Harrison and colleagues Figure 7b) comes from measurements by Girshick and colleagues^8^ over 6 levels of image resolution. It is unclear which resolution the green line is based on or why.

e None of the various decoding metrics (accuracy, precision, bias) in Harrison and collegues^1^ are specific to the commonly used inverted encoding model (IEM) in their paper. To illustrate this, we use a Mahalanobis distance decoder^15^ that yields qualitatively similar results as the IEM decoder (Supplemental Figure 1). We made other minor analysis changes to improve consistency and robustness (see Supplemental Methods), such as using channels *and* time-points as features for decoding, wider orientation bins, using repeated stratified random folds to split both real and simulated data, etc. (see Supplemental Methods).

f The same is true for the eccentricity of dot stimuli in the dataset from Bae^24^, shown as the inner ring in Figure 3a (*bottom*), which is close to edge of the full-field gratings used in Wolff and collegues^19^ where cardinal anisotropies are also observed (Figure 2a).

## References

1. Harrison, W. J., Bays, P. M. & Rideaux, R. Neural tuning instantiates prior expectations in the human visual system. Nat. Commun. 14, 5320 (2023).

2. Roth, M. M., Helmchen, F. & Kampa, B. M. Distinct Functional Properties of Primary and Posteromedial Visual Area of Mouse Neocortex. J. Neurosci. 32, 9716–9726 (2012).

3. Wang, G., Ding, S. & Yunokuchi, K. Difference in the representation of cardinal and oblique contours in cat visual cortex. Neurosci. Lett. 338, 77–81 (2003).

4. Fang, C., Cai, X. & Lu, H. D. Orientation anisotropies in macaque visual areas. Proc. Natl. Acad. Sci. 119, e2113407119 (2022).

5. Berman, N. E., Wilkes, M. E. & Payne, B. R. Organization of orientation and direction selectivity in areas 17 and 18 of cat cerebral cortex. J. Neurophysiol. 58, 676–699 (1987).

6. Dragoi, V., Turcu, C. M. & Sur, M. Stability of Cortical Responses and the Statistics of Natural Scenes. Neuron 32, 1181–1192 (2001).

7. Hubel, D. H. & Wiesel, T. N. Receptive fields, binocular interaction and functional architecture in the cat’s visual cortex. J. Physiol. 160, 106–154 (1962).

8. Blakemore, C. & Cooper, G. F. Development of the Brain depends on the Visual Environment. Nature 228, 477–478 (1970).

9. Hirsch, H. V. B. & Spinelli, D. N. Visual Experience Modifies Distribution of Horizontally and Vertically Oriented Receptive Fields in Cats. Science 168, 869–871 (1970).

10. Sengpiel, F., Stawinski, P. & Bonhoeffer, T. Influence of experience on orientation maps in cat visual cortex. Nat. Neurosci. 2, 727–732 (1999).

11. Liu, T., Cable, D. & Gardner, J. L. Inverted encoding models of human population response conflate noise and neural tuning width. J. Neurosci. 2453–17 (2017) doi:10.1523/JNEUROSCI.2453-17.2017.

12. Sprague, T. C. et al. Inverted Encoding Models Assay Population-Level Stimulus Representations, Not Single-Unit Neural Tuning. eNeuro ENEURO.0098-18.2018 (2018) doi:10.1523/ENEURO.0098-18.2018.

13. Rideaux, R., West, R. K., Rangelov, D. & Mattingley, J. B. Distinct early and late neural mechanisms regulate feature-specific sensory adaptation in the human visual system. Proc. Natl. Acad. Sci. 120, e2216192120 (2023).

14. Wei, X.-X. & Stocker, A. A. A Bayesian observer model constrained by efficient coding can explain ‘anti-Bayesian’ percepts. Nat. Neurosci. 18, 1509–1517 (2015).

15. Sprague, T. C., Boynton, G. M. & Serences, J. T. The Importance of Considering Model Choices When Interpreting Results in Computational Neuroimaging. eNeuro 6, (2019).

16. Gardner, J. L. & Liu, T. Inverted Encoding Models Reconstruct an Arbitrary Model Response, Not the Stimulus. eNeuro 6, (2019).

17. Wolff, M. J., Ding, J., Myers, N. E. & Stokes, M. G. Revealing hidden states in visual working memory using electroencephalography. Front. Syst. Neurosci. 9, (2015).

18. Wolff, M. J., Jochim, J., Akyürek, E. G. & Stokes, M. G. Dynamic hidden states underlying working-memory-guided behavior. Nat. Neurosci. 20, 864–871 (2017).

19. Wolff, M. J., Jochim, J., Akyürek, E. G., Buschman, T. J. & Stokes, M. G. Drifting codes within a stable coding scheme for working memory. PLOS Biol. 18, e3000625 (2020).

20. Himmelberg, M. M., Winawer, J. & Carrasco, M. Polar angle asymmetries in visual perception and neural architecture. Trends Neurosci. 46, 445–458 (2023).

21. Himmelberg, M. M., Winawer, J. & Carrasco, M. Linking individual differences in human primary visual cortex to contrast sensitivity around the visual field. Nat. Commun. 13, 3309 (2022).

22. Wandell, B. A., Dumoulin, S. O. & Brewer, A. A. Visual Field Maps in Human Cortex. Neuron 56, 366–383 (2007).

23. Bae, G.-Y. Neural evidence for categorical biases in location and orientation representations in a working memory task. NeuroImage 240, 118366 (2021).

24. Foster, J. J., Sutterer, D. W., Serences, J. T., Vogel, E. K. & Awh, E. The topography of alpha-band activity tracks the content of spatial working memory. J. Neurophysiol. 115, 168–177 (2015).

25. Foster, J. J., Bsales, E. M., Jaffe, R. J. & Awh, E. Alpha-Band Activity Reveals Spontaneous Representations of Spatial Position in Visual Working Memory. Curr. Biol. 27, 3216–3223.e6 (2017).

26. Carlson, T. A. Orientation Decoding in Human Visual Cortex: New Insights from an Unbiased Perspective. J. Neurosci. 34, 8373–8383 (2014).

27. Roth, Z. N., Heeger, D. J. & Merriam, E. P. Stimulus vignetting and orientation selectivity in human visual cortex. eLife 7, e37241 (2018).

28. Bae, G.-Y. & Luck, S. J. Dissociable Decoding of Spatial Attention and Working Memory from EEG Oscillations and Sustained Potentials. J. Neurosci. 38, 409–422 (2018).

29. Maloney, R. T. & Clifford, C. W. G. Orientation anisotropies in human primary visual cortex depend on contrast. NeuroImage 119, 129–145 (2015).

30. Freeman, J., Heeger, D. J. & Merriam, E. P. Coarse-Scale Biases for Spirals and Orientation in Human Visual Cortex. J. Neurosci. 33, 19695–19703 (2013).

31. Lennie, P. Distortions of Perceived Orientation. Nature. New Biol. 233, 155–156 (1971).

## Supplemental References

32. Ledoit, O. & Wolf, M. Honey, I shrunk the sample covariance matrix. J. Portf. Manag. 30, 110–119 (2004).

33. Myers, N. E. et al. Testing sensory evidence against mnemonic templates. eLife 4, e09000 (2015).

